# Research Productivity and Training Support for Doctoral Students in the Biological and Biomedical Sciences

**DOI:** 10.1101/2022.06.02.494498

**Authors:** Michael D. Schaller

## Abstract

Training of doctoral students as part of the next generation of the biomedical workforce is essential for sustaining the scientific enterprise in the US. Training of doctoral students primarily occurs at institutions of higher education and these trainees comprise an important part of the workforce at these institutions. Federal investment in the support of doctoral students in the biological and biomedical sciences is distributed differently than the distribution of students across different types of institutions, e.g. Public vs Private. Institutions in states that historically receive less federal support for research also receive less support for doctoral student training. Analysis of F31 awardees at different types of institution reveal little difference in research productivity and subsequent receipt of additional NIH awards. Analysis of a large cohort of doctoral students at different types of institutions also reveal little difference in research productivity, with the exception of citations. Thus, training outcomes, which are related to the quality of the student and training environment, are similar across different institutions. Factors that correlate with F31 funding include R01 funding levels and program size. The findings suggest strategies for institutions to increase success at securing F31s and modification of policy to promote more equitable distribution of F31s across institutions.

## Introduction

Long-term investment in biomedical research resulted in the rise of US research to a preeminent position in the world and the development of the knowledge base to combat disease and save lives. One recent illustration is the investment in basic research into mRNA vaccines, which led to the remarkably rapid development of highly effective vaccines against SARS-CoV2 during the pandemic (1, 2). During the years of a flat NIH budget, concerns about sustaining the biomedical research enterprise re-emerged amid concerns that the standing of US biomedical research in the world would erode and the development of medicines to manage disease would be impaired. Recommendations to sustain the enterprise focused on sustained and predictable funding for research, reduction of regulatory burden, modifying the biomedical workforce and modifying workforce training (3–7). The focus of this study relates to the biomedical workforce.

The estimated size of the US biomedical workforce between 2010 and 2014 was 289,147 to 305,500 persons (8, 9). Fifty percent of these employees worked in government and industry (9). Sixty percent of the workforce had an MS degree and between 30 and 46 % of the workforce held PhDs (8, 9). The estimated workforce supported by NIH extramural programs in FY09 was 121,465 full time effort positions, which provided support to 247,457 individuals (10). Trainees comprised 20% of the individuals supported by the NIH (10).

Advanced training of the biomedical workforce occurs predominantly in academic institutions of higher education with approximately 70% of PhD students training at Public institutions. The primary sources of support for training are fellowships, traineeships, research assistantships and teaching assistantships. PhD recipients can continue training as postdocs in a broad range of academic, industry and government positions, which can also be supported by fellowships, traineeships and research assistantships. The NIH and other federal sources provide significant resources for doctoral and postdoc training in the form of fellowships and institutional training grants, which provide traineeships to individuals. In FY17 the NIH investment in training grants was $1.5 billion, which was 4.5% of the NIH budget (11). The NIH F series of grants provide individual fellowships, the T series of grants provide institutional training grants and the K series of grants provide support for mentored career development. Evaluation of these grant mechanisms suggest that they are effective and promote research productivity and career advancement to subsequent research funding. Support on a training grant or an individual NIH fellowship increases success at securing a K award (12). K awardees publish more papers than non-awardees and are more successful at securing NIH research grants (11–13). Similar analysis of F32 awardees, who are postdocs supported by individual fellowships from the NIH, show increased research productivity in number of publications and increased success at securing NIH research grants (14, 15).

There are a number of concerns related to workforce training. First, the number of PhDs awarded exceeds the number of traditional positions, i.e. academic faculty positions (4, 16–18). Despite the fact that PhDs are employed in many positions outside of academia, e.g. in government or industry (3, 4), in the biomedical sciences there is a surplus of PhDs relative to job openings across all sectors (8, 19). Second, since many doctorates work outside of academia, they may require additional skillsets, which are not currently part of their training program, for success. (4, 17, 18). Third, the length of time spent in training as a doctoral student and in postdoctoral training was identified as a concern since this delays career progression and reduces earnings (6, 20). Recommendations were made to reduce the number of trainees, modify curricula to prepare graduates for broader careers, shorten the time to degree and increase the transparency of outcomes of training programs. There have also been recommendations to shift training support to federal fellowships and training grants (6, 20). Review of the proposed training plan as part of the review of applications is expected to strengthen training. Further increased trainee success is expected, given the productivity and successful career progression of NIH supported trainees. While the merits of this recommendation are clear, careful consideration of its full potential impact is required. This study begins this assessment by examining the university biomedical workforce, how trainees (specifically doctoral students) are supported at different institutions and the research productivity and success of students at different institutions.

## Methods

### Sources of Data

NIH award data and publication data associated with NIH awards was captured from NIH Reporter (https://reporter.nih.gov/). Information about trainees and the workforce at academic institutions is from the Graduate Students and Postdoctorates in Science and Engineering Survey (GSS) (https://www.nsf.gov/statistics/srvygradpostdoc/). The data tables for each year from 1985 through 2019 was used for this analysis (21). Data regarding doctorates awarded is from the Survey of Earned Doctorates (SED) (https://www.nsf.gov/statistics/srvydoctorates/). The data tables for each year from 1994 through 2020 were used for this analysis (22). GSS and SED searches for data on numbers of doctorates, postdocs and sources of support were restricted to the field of study of Biological and Biomedical Sciences. Information about doctoral dissertations was retrieved from the ProQuest Dissertations & Theses Global database. PubMed was searched for publication data using the BioEntrez package from Python (23). Data on research expenditures was from the NCSES National Patterns of R&D Resources (24)(https://www.nsf.gov/statistics/natlpatterns/).

### Cohorts

Data was collected on two cohorts of students, 1) F31 awardees from 2001 to 2016 inclusive, and 2) all doctorates completing their dissertation from 2012 through 2016. For the latter cohort, doctorates and their advisors were identified from the ProQuest Dissertations & Theses Global database (www.proquest.com – access provided by West Virginia University Libraries). The database was searched for institutions designated as doctorate-granting institutions by GSS (235 institutions), and the search limited to subjects listed as Biological and Biomedical Sciences in the GSS. This provided a list of doctorates and advisors for each of 235 institutions. Doctorates with no affiliated advisor listed were excluded from the analysis (694 doctorates).

### Publications and citations

F31 awardee publication data was extracted from NIH Reporter in April and May 2021. The data was curated to ensure all publications include the awardee as an author and to correct misspellings of awardee names in the author list. Curation impacted a few percent of all publications listed. The number of first author publications and total publications for each F31 awardee were calculated using a Python script. Publications by the doctorates identified from the ProQuest database were found by searching PubMed using the Python BioEntrez package (23). Search criteria for first author publications included the doctorate’s name as first author, the advisor’s name as an author and the institution name as affiliation. The search for first author publications was performed in May and June 2021. Search criteria for total publications included the doctorate’s and advisor’s names and the institution name as affiliation. The search for total publications was performed in November and December 2021. In many cases, more than one advisor was listed per doctorate. PubMed searches using each advisor as a co-author with the doctorate were performed to identify all publications associated with the doctoral candidate. The year of each publication was also captured. Citations for both cohorts were counted using the BioEntrez package for Python and the search was performed in November 2021 (23). PMIDs were used to search PubMed for the list of PMCIDs that cite each publication and the number of PMCIDs counted.

### Grant success by F31 awardees

F31 awardees successfully securing additional NIH funding were identified by matching NIH PI IDs between F31 awards and other NIH awards including F32, K99, R15, R21 and R01 awards through fiscal year 2021.

### Institutional Review Board

The West Virginia University IRB approved the study. IRB approval numbers are WVU Protocol#: 2202521185 and WVU Protocol#: 2203537777.

### Statistics

None of the data exhibit a Gaussian distribution, therefore nonparametric statistics were used for the analysis. Some of the data presented in the tables cannot be directly compared statistically, since the groups are not independent. Statistical comparisons were made between independent groups, i.e. Public vs Private Institutions, Land Grant vs non-Land Grant institutions and IDeA vs non-IDeA institutions. Publication and citation data was compared using the Mann Whitney test. Rates of securing additional NIH funding data was compared using Fisher’s Exact test. Details of the statistical comparisons are presented in Supplemental Table 1.

## Results

According to the National Center for Science and Engineering Statistics, research investment in 2019 totaled approximately $656 billion, of which $107 billion was invested in basic research (24). In 2019, 46% of basic research expenditures occurred at higher education institutions (24). In addition to serving as performance sites for biomedical research, institutions of higher education are also the primary training centers for the biomedical workforce. Consequently, a large percentage of the biomedical workforce at these institutions is comprised of trainees. Thus, the investment in trainees is a critical investment in the future of biomedical research and in current basic research, which is the foundation for tomorrow’s cures.

In the biological and biomedical sciences alone, the workforce at higher educational institutions in 2019 was estimated to include 45,466 doctoral students and 19,631 postdoctoral fellows (25). The balance of the workforce between students and postdocs varies between institutions. At public institutions, approximately 68% of the workforce is comprised of doctoral students, whereas at private institutions, ∼55% of the workforce are doctoral students (Fig. 1). The workforce was also examined at two other classifications of university, land grant and IDeA institutions. Land grant institutions were originally designated by states for donations of federal land or money to establish the university and frequently include making contributions to benefit the state as part of their mission. The Institutional Development Award (IDeA) program was congressionally mandated in 1992 as a mechanism to provide investment to build research infrastructure in states with historically low levels of funding from the NIH. The doctoral student component of the workforce at land grand institutions is similar to that of public institutions. IDeA institutions have the largest percentage of the workforce comprised of graduate students (∼72%). Approximately 10% of the workforce at each type of institution consists of non-faculty researchers and the postdoctoral component of the workforce ranges from 18% (IDeA institutions) to 34% (private institutions). As a result of the difference compositions of the workforce, policies affecting trainees will have different impacts at different institutions.

**Figure 1.**
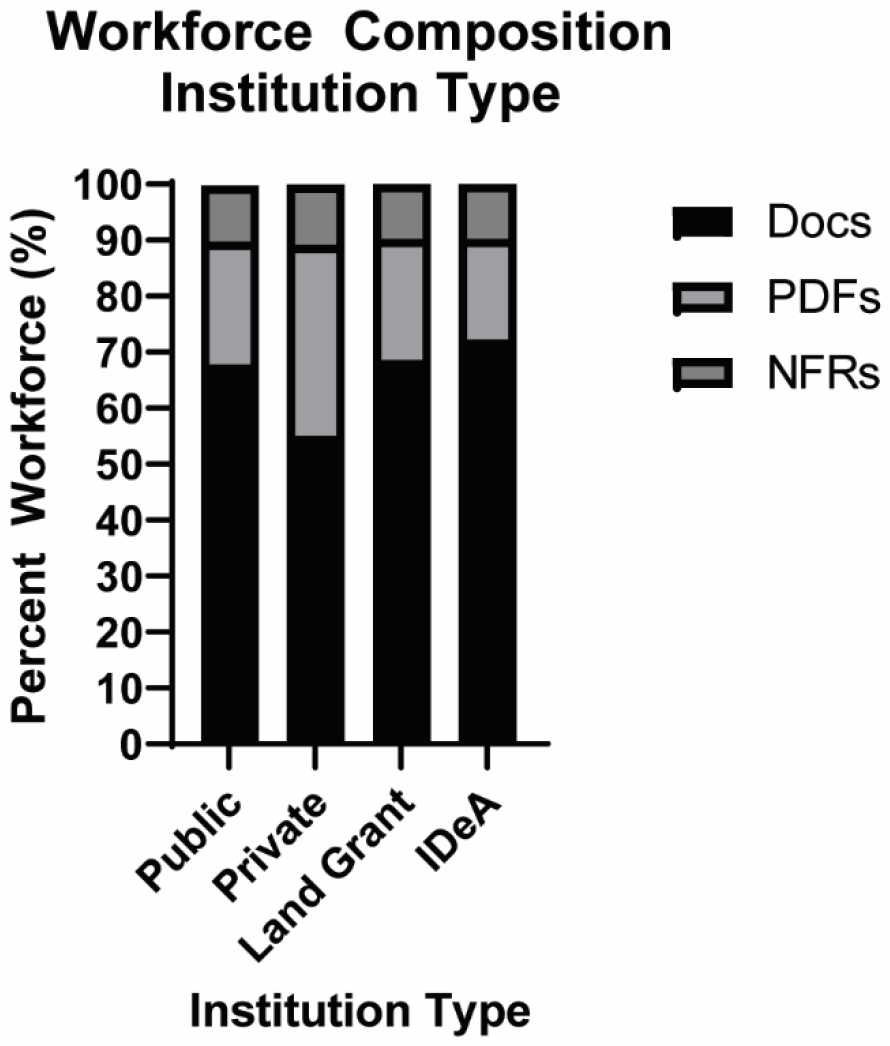
Composition of the workforce at different types of institutions. The GSS reports the number of doctoral students (Docs), postdocs (PDFs) and non-faculty researchers (NFRs) affiliated with different programs at each university. The number of individuals in each of these categories, in biological and biomedical sciences programs in 2019 were tallied for all Private, Public, Land Grant and IDeA institutions and the percentage of individuals in each category at each type of institution was calculated.

### Distribution of Trainees

Approximately 65% of biological and biomedical sciences doctoral students trained at public universities and 35% trained at private institutions in 2019 (Table 1)(25). The data is consistent between the Graduate Students and Postdoctorates in Science and Engineering Survey (GSS) and the Survey of Earned Doctorates (SED) (Table 1) (25, 26). Land Grant institutions trained between 25% and 30% of doctoral students. Institutions in IDeA states trained approximately 11% of doctoral students in 2019. In 2019, approximately 50% of postdocs in academic institutions trained at Public institutions and 50% trained at Private institutions (Table 1). Land Grant institutions trained 22% of postdocs and 6% of postdocs trained at IDeA institutions.

**Table 1.** Number of Trainees in 2019 (Biological and Biomedical Sciences Trainees)

| **Institution Type** | **# Doctoral Students (GSS)** | **% Doctoral Students (GSS)** | **# Doctorates Awarded (SED)** | **% Doctorates Awarded (SED)** | **# Postdocs Trained (GSS)** | **% Postdocs Trained (GSS)** |
| --- | --- | --- | --- | --- | --- | --- |
| **Public** | 29,371 | 64.6% | 5,594 | 64.3% | 9,629 | 49.1% |
| **Private** | 16,095 | 35.4% | 2,785 | 32.0% | 10,002 | 51.0% |
| **Land Grant** | 13,461 | 29.6% | 2,156 | 24.8% | 4,246 | 21.6% |
| **IDeA** | 4,829 | 10.6% | 988 | 10.9% | 1,204 | 6.1% |
| **All** | 45,466 |  | 8,702 |  | 19,631 |  |

Data on the distribution of doctoral students and postdocs amongst different types of institutions (from the GSS) and the number of doctorates awarded by different types of institutions (from the SED).

The SED provides a historical record of the number of doctorates awarded (the GSS only parses graduate students into masters and doctoral students since 2017). The number of biological and biomedical doctorates awarded has plateaued since 2009 and has declined since 2015 (Fig 2A). Simple linear regression of the data from 2009 to 2020 indicates an R squared value of 0.2167 and that the slope does not significantly differ from zero (Supp Fig 1A). Analysis of the data from 2015 to 2020 reveals an R squared value of 0.5721 and a negative slope that differs from zero (p value = 0.049) (Supp Fig 1B). This trend is encouraging given longstanding concerns about the large numbers of students earning PhDs each year (4, 16–18). The percentage of students earning doctoral degrees from public, private, land grant and IDeA institutions has been relatively constant since 1994 (Fig 2B). The number of postdocs in the biological and biomedical sciences appears to have peaked in 2010, declined and plateaued since then (Fig 3A). The percentage of postdocs trained at Public, Private, Land Grant and IDeA institutions has been relatively constant since 2010 (Fig 3B).

**Figure 2.**
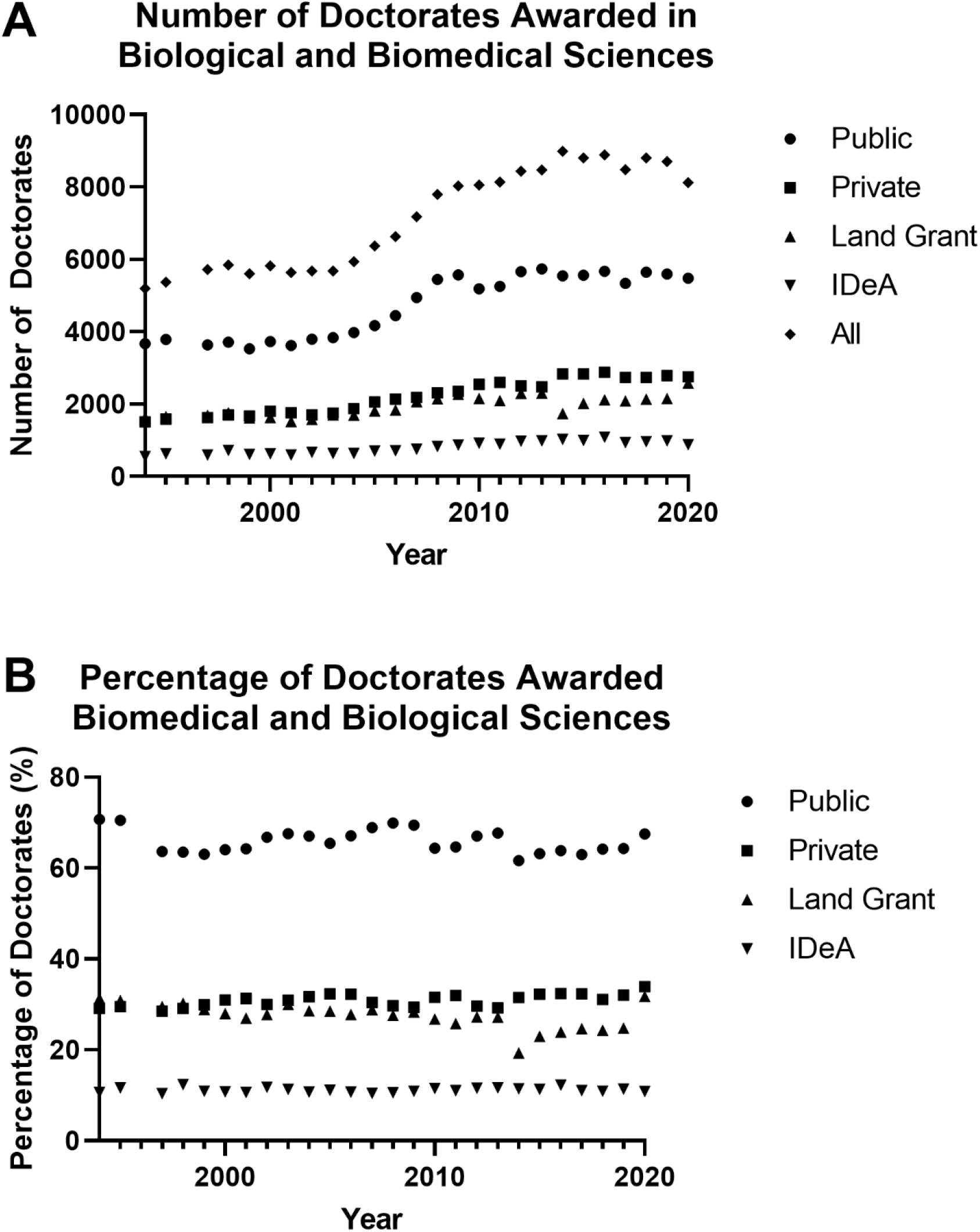
Historical record of doctorates awarded in biological and biomedical sciences at US Universities. The number of doctorates awarded in biological and biomedical sciences from 1994 to 2020 was retrieved from the SED (22). The number of doctorates awarded (**panel A**) and percentage of total doctorates awarded (**panel B**) at Private, Public, Land Grant and IDeA institutions was calculated.

**Figure 3.**
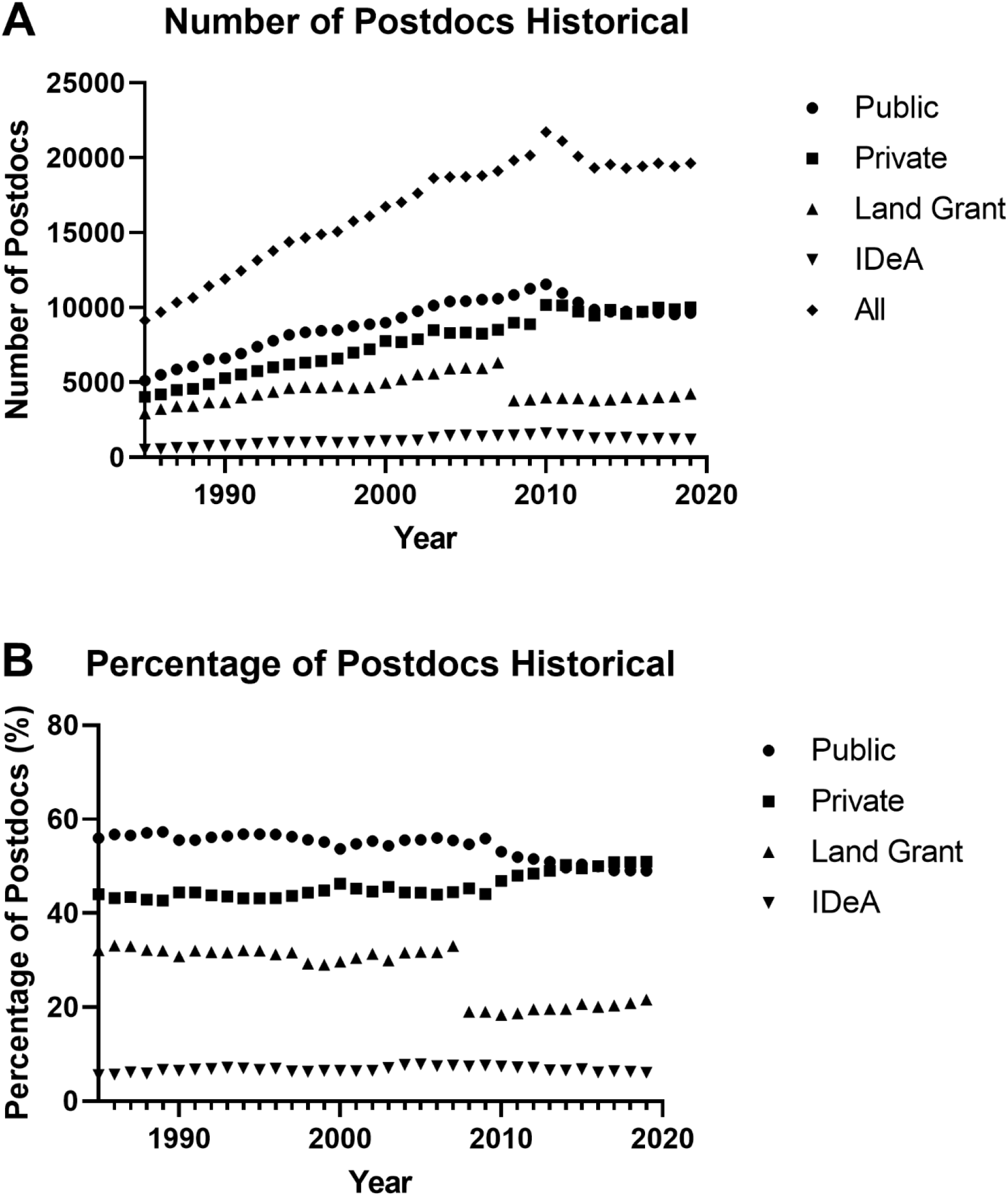
Historical record of the number of postdocs in biological and biomedical sciences at US universities. The number of postdocs working in biological and biomedical sciences from 1985 to 2019 was retrieved from the GSS (21). The number of postdocs (**panel A**) and percentage of total postdocs (**panel B**) at Private, Public, Land Grant and IDeA institutions was calculated.

### Trainee Support

The major mechanisms of financial support of doctoral students are fellowships, traineeships, research assistantships and teaching assistantships (25). In 2019, 12.52% of all doctoral students were supported on federal fellowships and traineeships. Excluding non-training grant eligible students, approximately 24.67% of doctoral students were supported on federal fellowships and traineeships. A larger percentage of students at private institutions are supported on fellowships and traineeships than students at other types of institutions (Fig 4A). Conversely, the percentage of students supported by research assistantships and teaching assistantships at public, land grant and IDeA institutions is greater than the percentage of students supported by similar mechanisms at private institutions (Fig 4A). Nearly 50% of doctoral students at Public, Land Grant and IDeA institutions are supported by research assistantships and approximately 20% are supported by teaching assistantships. The federal government provides critical support for doctoral student training. In 2019, federal funds supported 29% of students supported by fellowships, 60.5% of the students supported by traineeships and 47.4% of research assistantships (25). Institutions supported 97.3% of teaching assistantships (25). The percentage of doctoral students receiving federal support at different types of institutions is shown in Fig 4B. The highest percentage of students receiving federal fellowships are at Private and Land Grant Institutions. The percentage of students supported on federal traineeships at Private institutions is more than twice as high as at any other type of institution. The percentage of students at IDeA institutions supported by federal fellowships and traineeships lags behind all the other types of institutions. The percentage of students supported by federal research assistantships at different types of institutions is more comparable. The major source of federal funding for biomedical research is the NIH. Support for trainees on NIH fellowships, traineeships and research assistantships is illustrated in Fig 4C. Since NIH is the major source of funding in the area, the trends parallel the trends seen in total federal funding. Interestingly, the highest percentage of students supported by NIH research assistantships are at Private institutions.

**Figure 4.**
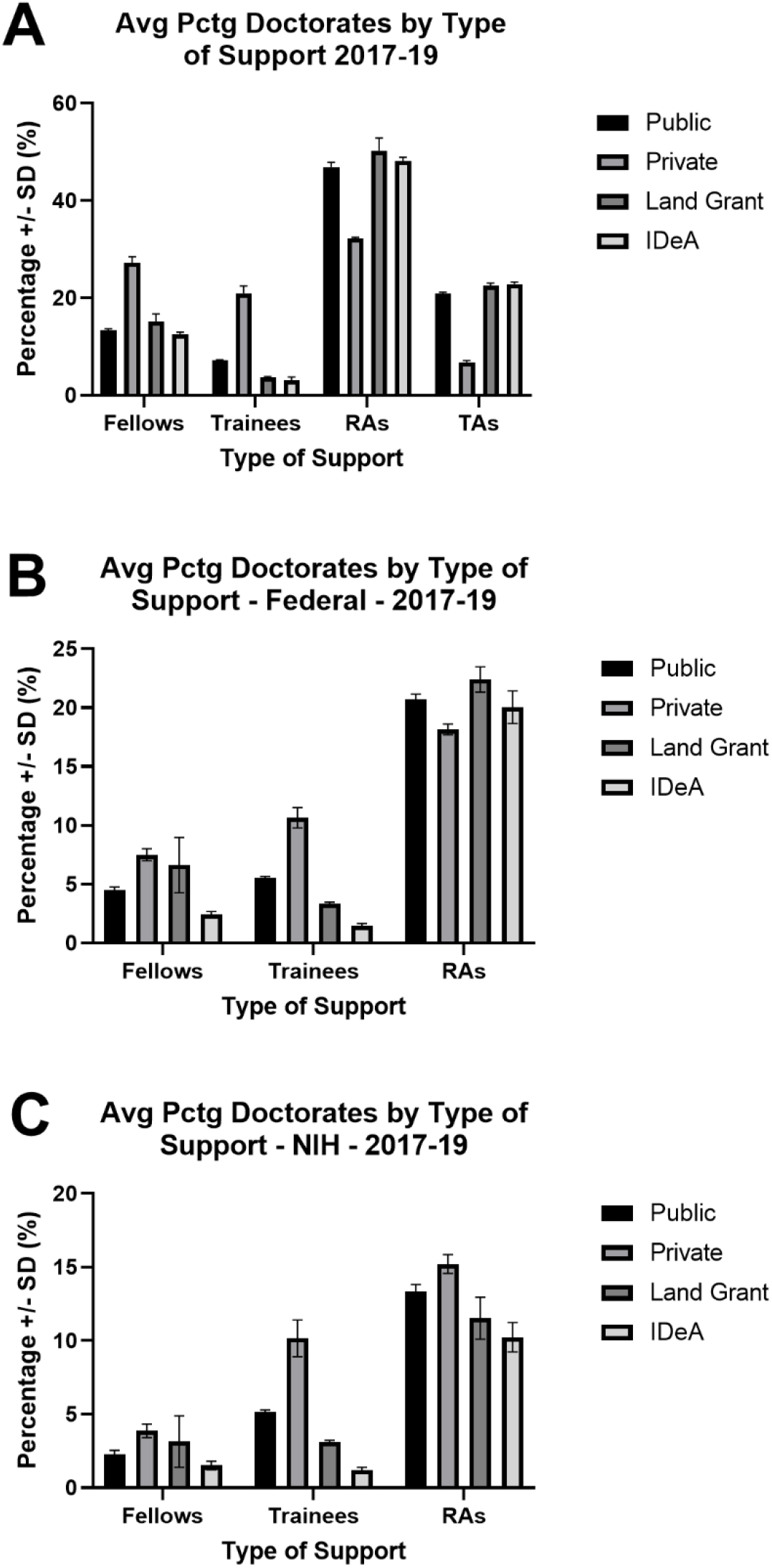
Source of financial support for doctoral students in biological and biomedical sciences. The source of financial support for doctoral students from 2017 to 2019 was retrieved from the GSS (21). The percentage of students supported by different mechanisms at Public, Private, Land Grant and IDeA institutions is shown. **A)** Support from all sources. **B)** Support from federal sources. **C)** Support from the NIH. Fellows = fellowships, Trainees = traineeships, RAs = research assistantships, TAs = teaching assistantships. Note that federal sources do not support teaching assistantships.

The major sources of support for postdocs are fellowships, traineeships and research grants. The majority of postdocs at all types of institutions are supported on research grants (65% to 68%) (Fig 5). In general, the proportion of postdocs on fellowships and traineeships were comparable at Private and Public institutions and the percentage of postdocs supported by fellowships and traineeships at Land Grant and IDeA institutions lagged (Fig 5).

**Figure 5.**
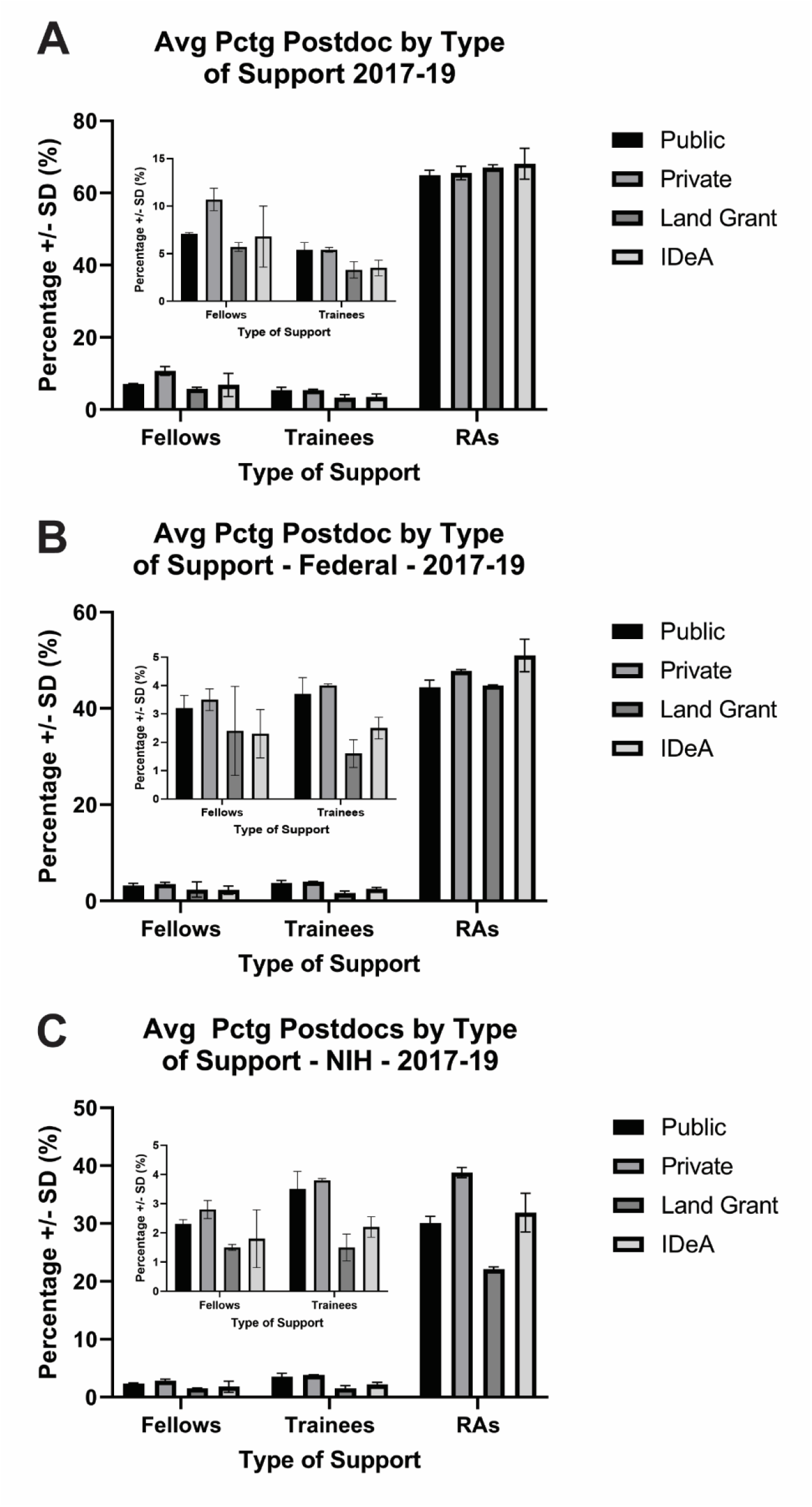
Source of financial support for postdocs in biological and biomedical sciences at US universities. The source of financial support for postdocs from 2017 to 2019 was retrieved from the GSS (21). The percentage of postdocs supported by different mechanisms at Public, Private, Land Grant and IDeA institutions is shown. **A)** Support from all sources. **B)** Support from federal sources. **C)** Support from the NIH. Fellows = fellowships, Trainees = traineeships, RAs = research assistantships. The inset in each panel shows the data for fellowships and traineeships with an appropriate y-axis scale to better evaluate differences.

### Trainee Support from the NIH

A history of NIH support for doctoral student fellowships (F31 awards) is illustrated in Fig 6A. For the last twenty years there was a sustained increase in the number of F31 awards. The percentage of F31 awards made to different types of institutions is illustrated in Fig 6B. The striking observation from this analysis is that approximately 50% of F31s are awarded to students at Private institutions and to students at Public institutions. This is in contrast to the observations that 65% of doctoral students train at Public institutions and 35% train at Private institutions (Figure 2). The percentage of F31s awarded to Land Grant and IDeA institutions are about half of the percentage of doctoral students that train at those institutions.

**Figure 6.**
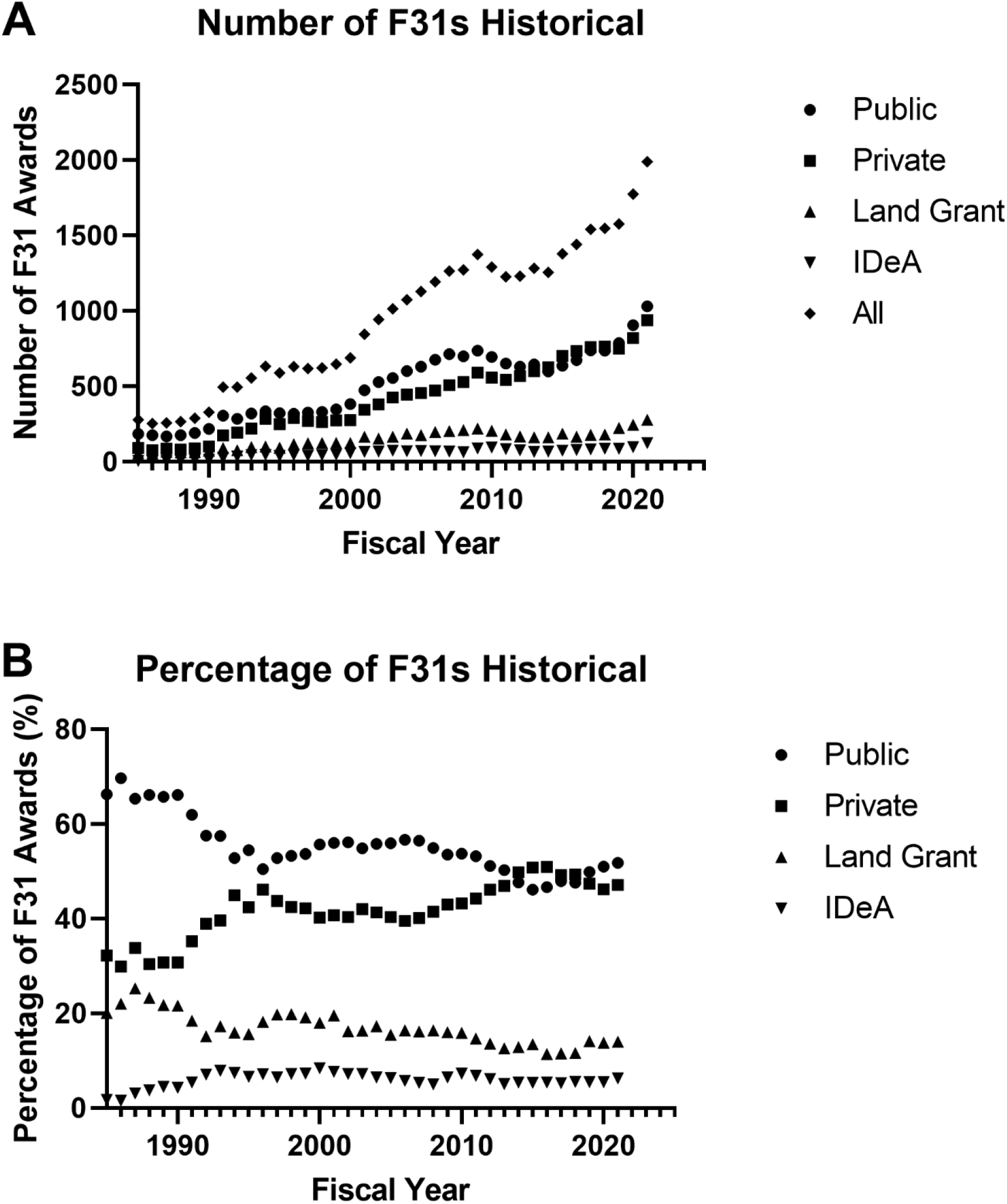
Historical record of F31 awardees. The number of F31s for each fiscal year from 1985 to 2021 was tallied using data acquired using NIH Reporter. The number of awards (**panel A**) and the percentage of awards (**panel B**) at Public, Private, Land Grant and IDeA institutions is plotted.

The historical record of NIH postdoc fellowships (F32 awards) is shown in Figure 7. The number of F32 awards has declined since the mid-1990s. Since 2007, K99 awards from the NIH also supported postdocs, but the number of awards is small compared with the F32 program. The percentage of F32 awardees at Private institutions was higher than the percentage at Public institutions but was comparable in the most recent fiscal year (FY21). Note that F32 awardees at research institutions and in industry were not captured as part of the analysis. The percentage of F32 awardees at Land Grant and IDeA institutions was approximately half of the percentage of postdocs trained at these institutions. One other mechanism for the support of trainees by NIH is the T32 training grant. The number of T32s has declined and plateaued since 2010 (Fig 8). Approximately 50% of T32s were held at Public and 50% at Private institutions. Land Grant institutions held approximately 13% of T32s and IDeA institutions were awarded approximately 5% of T32s. This analysis does not address the number of trainees supported by T32 training grants, since multiple trainees are concurrently supported by the same T32 and there is a wide range in the number of trainees supported by each award. Further, some T32s support doctoral students, some T32s support postdocs and some T32s support both. Therefore, the analysis of T32 support is much more complex than the analysis of fellowships and will not be addressed further.

**Figure 7.**
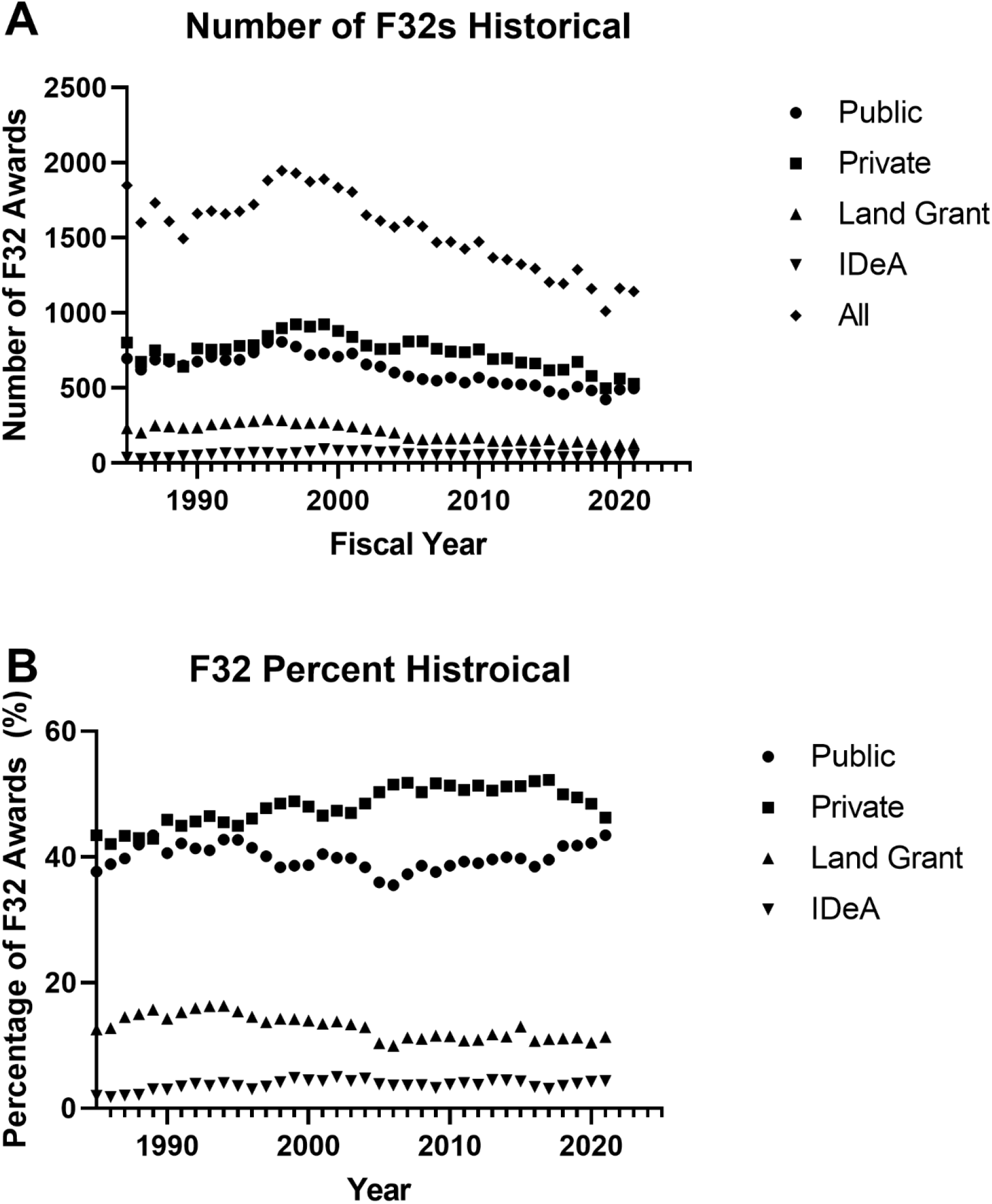
Historical record of F32 awardees. The number of F32s for each fiscal year from 1985 to 2021 was tallied using data acquired using NIH Reporter. The number of awards (**panel A**) and the percentage of awards (**panel B**) at Public, Private, Land Grant and IDeA institutions is plotted.

**Figure 8.**
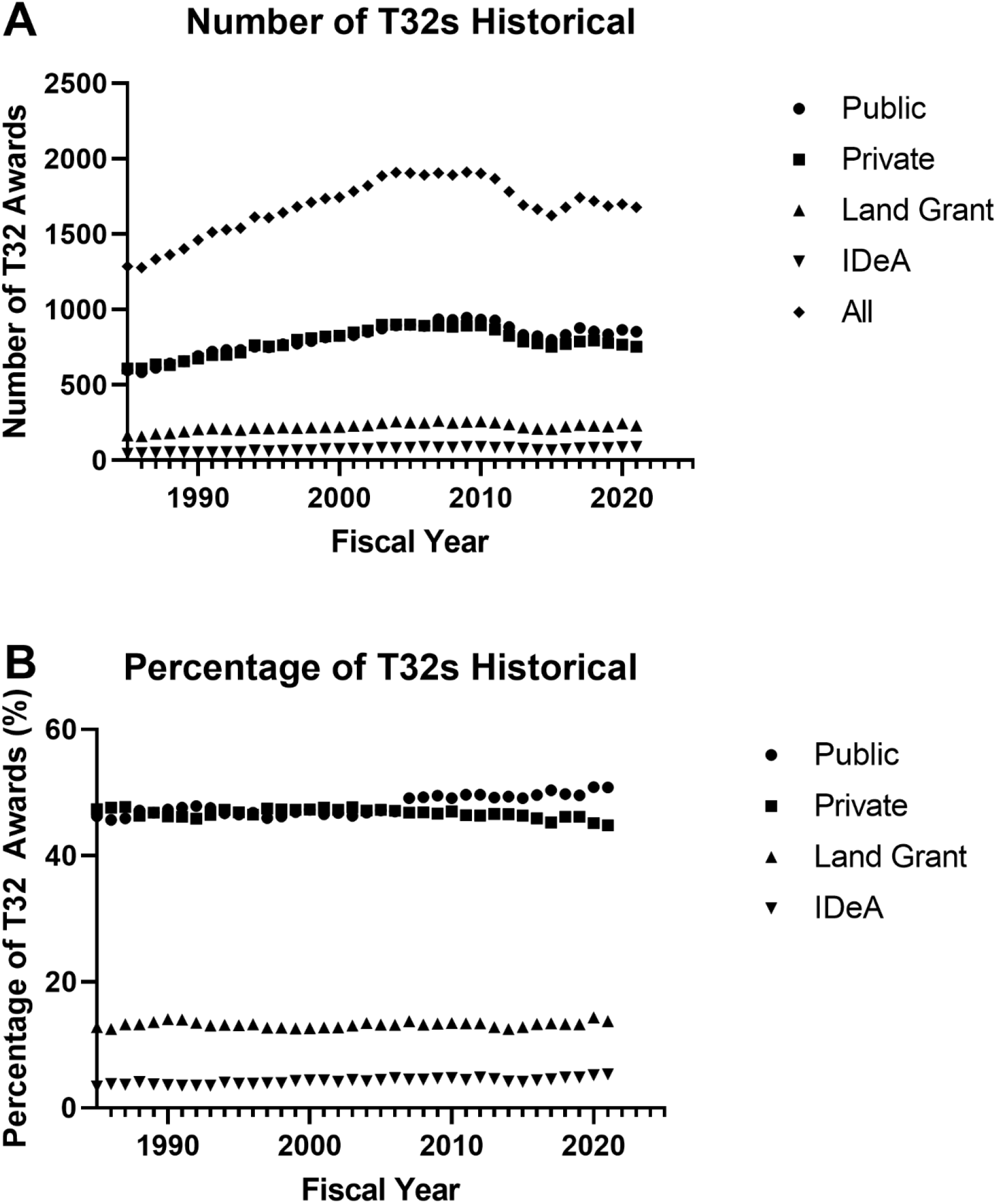
Historical record of T32 awardees. The number of T32s for each fiscal year from 1985 to 2021 was tallied using data acquired using NIH Reporter. The number of awards (**panel A**) and the percentage of awards (**panel B**) at Public, Private, Land Grant and IDeA institutions is plotted.

Analysis of trainee support by NIH shows that postdoc support at Public and Private institutions parallels the number of trainees at these institutions, while trainee support at Land Grant and IDeA institutions lag. Doctoral support at different types of institutions is not comparable to the number of trainees at these institutions. These observations raise questions about support for the training of doctoral students including the factors leading to successful competition for F31 awards, strategies to employ to increase F31 success rates and policies to support the training of the biomedical workforce of the future.

### Correlations with number of F31s awarded

#### NIH Funding

In FY19 there were 26,861 R01s administered across all NIH institutes. Of these, 23,616 were held at US academic institutions. The percentage of doctorates awarded, F31s held, and R01s held at different types of institutions is compared in Table 2. The percentage of F31s awarded at different types of institutions more closely reflects the percentage of R01s than the percentage of doctorates. Focusing only on institutions holding an F31 in FY19, the number of F31s held at different institutions in FY19 correlated with the number of R01s held at those institutions (Fig 9A). The number of F31s held at different institutions in FY19 also correlated with the number of doctorates awarded at those institutions in 2019 (Fig 9B).

**Figure 9.**
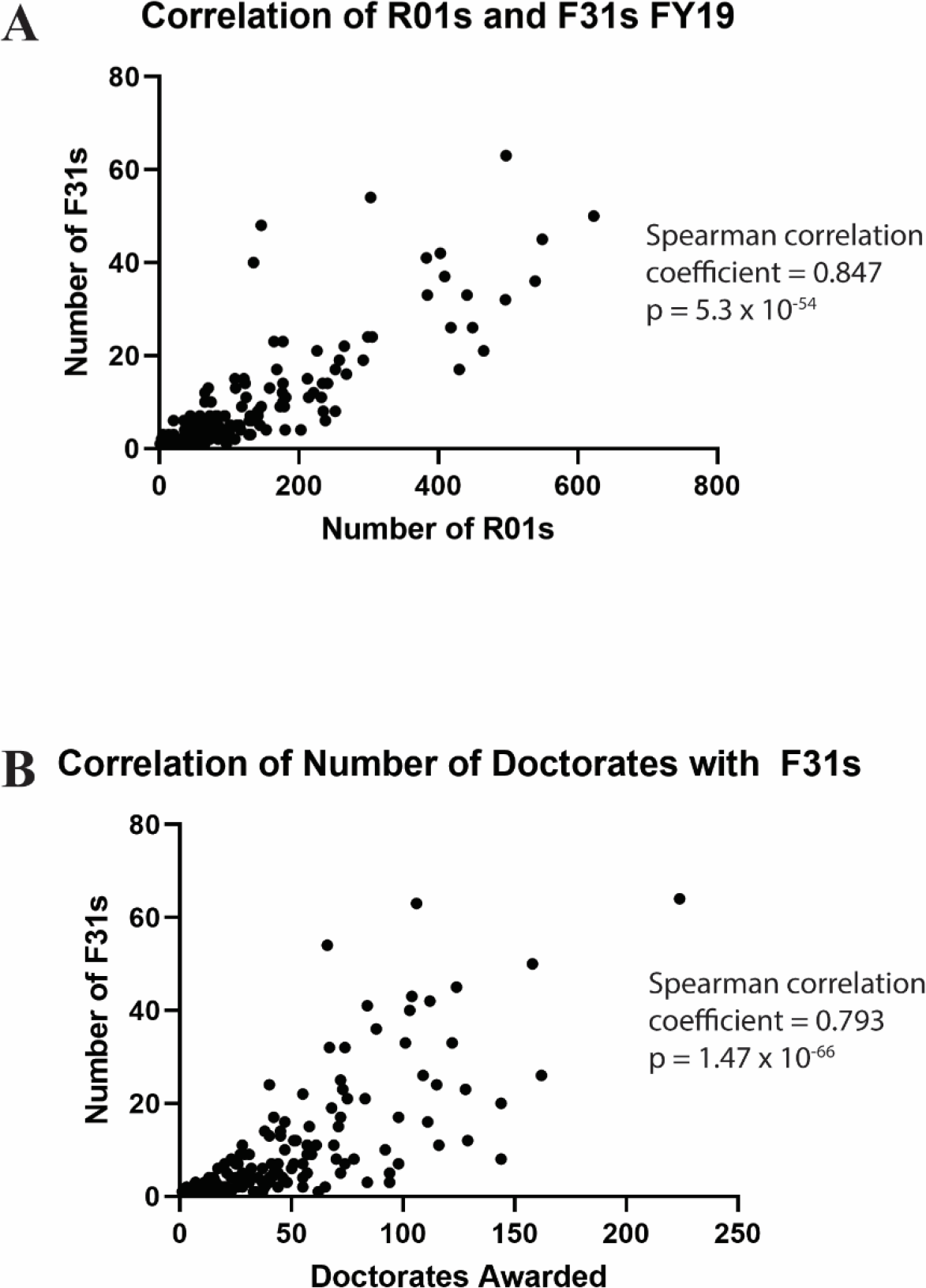
Correlations with number of F31 awards. **A)** The number of F31s at individual institutions is plotted against the number of R01s held at the institution during FY19. F31 and R01 data was extracted using NIH Reporter. **B)** The number of F31s at individual institutions during FY19 is plotted against with number of doctorates awarded at the institution in 2019. F31 data was calculated using data from NIH Reporter. The number of doctorates awarded is from the SED (22).

**Table 2.** Comparison of Earned Doctorates, F31s Awarded and R01s Awarded at Different Types of Institutions in FY19.

|  | **Public** | **Private** | **Land Grant** | **IDeA** |
| --- | --- | --- | --- | --- |
| Percent Doctorates 2019 | 64.3% | 32.0% | 24.8% | 11.4% |
| Percent F31s FY19 | 49.9% | 47.4% | 14.1% | 5.5% |
| Percent R01s FY19 | 53.9% | 46.1% | 15.4% | 6.7% |

The percentage of doctorates awarded is based upon data from the SED. The percentage of F31 and R01 awards was extracted from NIH data using NIH Reporter.

#### Training Grant Eligibility

The proportion of the doctoral student population that was training grant eligible (TGE) was estimated from the GSS by tallying the total number of US doctoral students and the total number of doctoral students. In 2019, 73.1% of biological and biomedical graduate students were training grant eligible. Private institutions had the highest percentage of TGE doctoral students (74.9%) followed by Public institutions (72.2%). Land Grant institutions (70.7%) and IDeA institutions (69.6%) had lower percentages of TGE doctoral students. Part of the difference in the number of F31 awards at different institutions can be attributed to the number of students who are training grant eligible.

#### Awardee Performance

Another variable in the selection of F31 awardees is the quality of applicant from different types of institutions. Quality was evaluated using training outcome as proxy. Outcome was measured as success in advancing on an independent research trajectory and research productivity. Research success was measured by securing additional funding from NIH such as an F32/K99 and R series grants. Research productivity was measured by the number of publications and citations. The cohort of all F31 awardees from all NIH Institutes from 2001 to 2016 inclusive was selected for this analysis. This cohort consisted of 4,579 F31 awardees from Public institutions, 3,714 from Private institutions, 1,323 from Land Grant institutions and 544 F31 recipients from IDeA institutions. The dates for inclusion in the cohort provided a large number of individual records for analysis and provided time for the cohort to complete training, finish publications, accumulate citations and secure additional funding. Note that a single awardee can appear in multiple categories, e.g. an awardee at an IDeA institution that is a Land Grant institution and a Public university.

Approximately 9 to 10% of this cohort of F31 awardees were also awarded an F32 and 2 to 3% were awarded a K99 (Table 3). Fisher’s exact test revealed no difference in the rate of success at securing a more advanced NIH fellowship between students holding F31s at different types of institutions. Approximately 6 to 7% of this cohort of F31 awardees were also awarded an R01 and 4 to 5% were awarded an R21 (Table 3). Fisher’s exact test revealed no difference in the rate of success at securing an R01 for F31 awardees at different types of institutions. There was a significant difference in the percentage of F31 awardees at Private and Public institutions who were also awarded an R21. A small number of awardees secured R15, R35, R41, R42, R43 and R44 grants. The small numbers precluded meaningful analysis. Using securing additional NIH funding as a metric of scientific success, this analysis demonstrates that F31 awardees from different types of institutions are comparably successful. A striking observation was where F31 awardees eventually held their R01s (Table 4). More than half of the F31 awardees at Private institutions who secured R01 funding held their R01s at Private institutions. Approximately 75% of F31 awardees at Public or Land Grant institutions, who successfully competed for R01 awards, held their R01s at Public institutions. Almost 40% of F31 awardees at IDeA institutions held their R01s at IDeA institutions. While there are multiple factors impacting the career trajectory of doctoral students, this observation suggests that the distribution of fellowships across different institutions might contribute to shaping the distribution of successful faculty in the future.

**Table 3.**
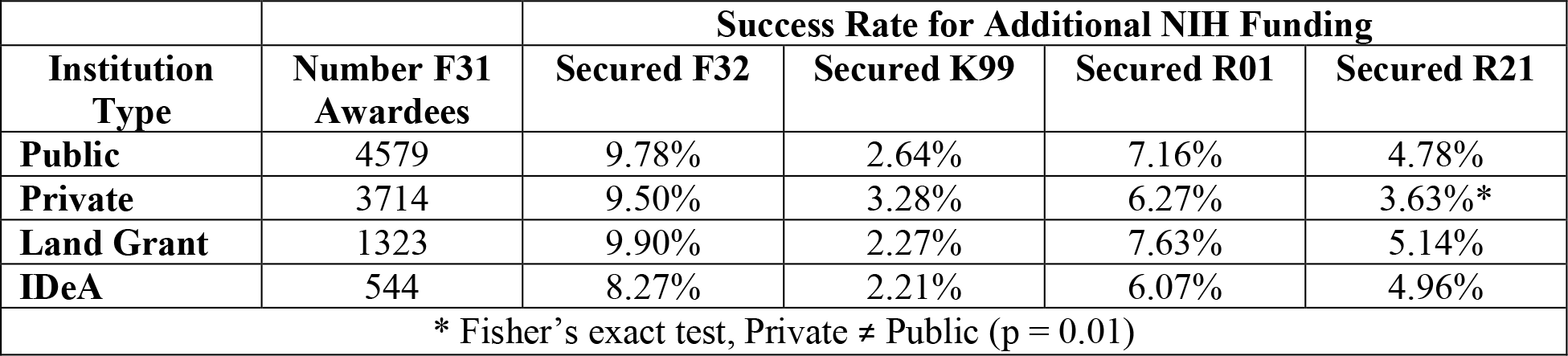
Percentage of F31 Awardees who secure additional Fellowships or R series Awards.

| **Institution Type** | **Number F31 Awardees** | **Success Rate for Additional NIH Funding** |  |  |  |
| --- | --- | --- | --- | --- | --- |
|  |  | **Secured F32** | **Secured K99** | **Secured R01** | **Secured R21** |
| **Public** | 4579 | 9.78% | 2.64% | 7.16% | 4.78% |
| **Private** | 3714 | 9.50% | 3.28% | 6.27% | 3.63% * |
| **Land Grant** | 1323 | 9.90% | 2.27% | 7.63% | 5.14% |
| **IDeA** | 544 | 8.27% | 2.21% | 6.07% | 4.96% |
| * Fisher's exact test, Private ≠ Public (p = 0.01) |  |  |  |  |  |

**Table 4.** Comparison of Types of Institutions Where F31 Awardees Hold R01s.

|  | **Public F31** | **Private F31** | **Land Grant F31** | **IDeA F31** |
| --- | --- | --- | --- | --- |
| **Public R01** | 74.1% | 34.3% | 75.2% | 69.7% |
| **Private R01** | 19.8% | 55.4% | 21.8% | 24.2% |
| **Land Grant R01** | 18.6% | 11.2% | 31.7% | 18.2% |
| **IDeA R01** | 13.7% | 5.6% | 16.8% | 39.4% |

The research productivity of F31 awardees at different institutions was also compared. First, the publication record of F31 awardees as listed in NIH REPORTER was compared. The average number of first author publications for the 8,293 F31 awardees in the cohort was 1.71 +/- 1.79 papers and the average number of total publications was 2.57 +/- 2.83 papers (see Tables 5 & 6). The average number of first author publications by F31 awardees at different types of institutions ranged from 1.58 +/- 1.61 (Private institutions) to 1.87 +/- 1.87 (Land Grant Institutions) papers. The average number of first author publications by F31 awardees at Public universities was greater than F31 awardees at Private institutions. The average number of first author publications of F31 awardees at Land Grant institutions was greater than F31 awardees at non-land grant universities. The median number of first author publications at each type of institution was the same (Table 5). The average number of total publications ranged from 2.47 +/- 2.6 (Private institutions) to 2.72 +/- 3.02 (Land Grant institutions) papers (Table 6). The differences in average number of total publications between F31 awardees at different types of institutions was not statistically different.

**Table 5.**
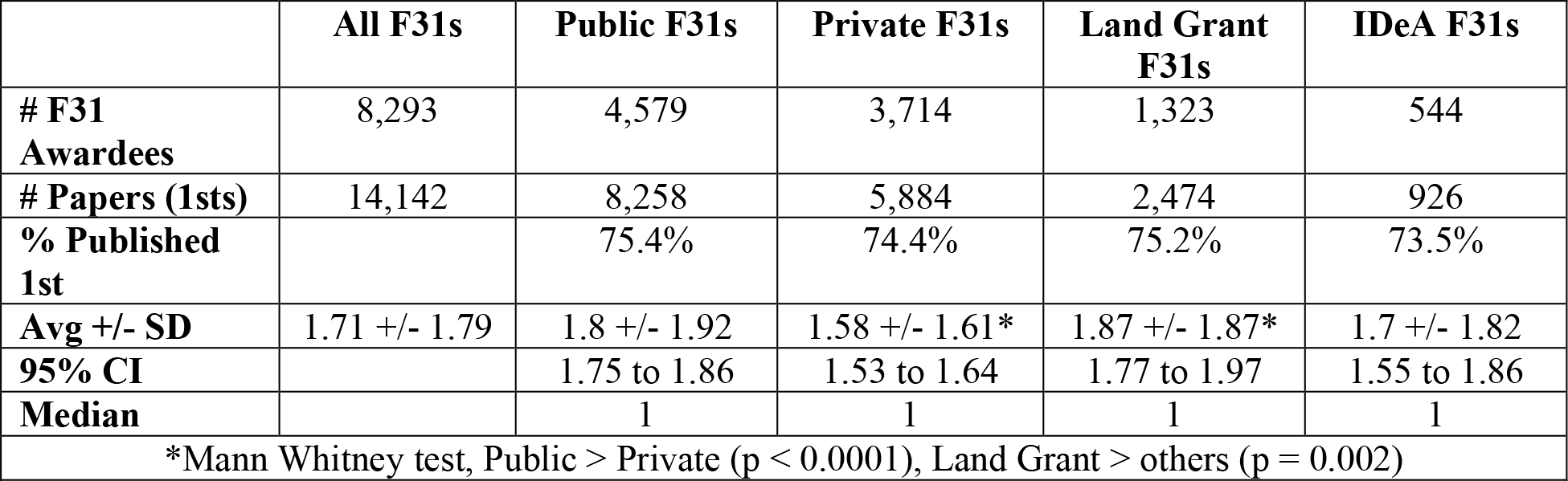
Comparing of Number of First Author Publications of F31 Awardees.

|  | **All F31s** | **Public F31s** | **Private F31s** | **Land Grant F31s** | **IDeA F31s** |
| --- | --- | --- | --- | --- | --- |
| **# F31 Awardees** | 8,293 | 4,579 | 3,714 | 1,323 | 544 |
| **# Papers (1sts)** | 14,142 | 8,258 | 5,884 | 2,474 | 926 |
| **% Published 1st** |  | 75.4% | 74.4% | 75.2% | 73.5% |
| **Avg +/- SD** | 1.71 +/- 1.79 | 1.8 +/- 1.92 | 1.58 +/- 1.61* | 1.87 +/- 1.87* | 1.7 +/- 1.82 |
| **95% CI** |  | 1.75 to 1.86 | 1.53 to 1.64 | 1.77 to 1.97 | 1.55 to 1.86 |
| **Median** |  | 1 | 1 | 1 | 1 |
| *Mann Whitney test, Public > Private (p < 0.0001), Land Grant > others (p = 0.002) |  |  |  |  |  |

**Table 6.** Comparison of Total Publications of F31 Awardees.

|  | **All F31s** | **Public F31s** | **Private F31s** | **Land Grant F31s** | **IDeA F31s** |
| --- | --- | --- | --- | --- | --- |
| **# F31 Awardees** | 8,293 | 4579 | 3714 | 1323 | 544 |
| **# Papers (Total)** | 21,318 | 12,147 | 9,171 | 3,599 | 1,424 |
| **% Published** |  | 80.0% | 79.9% | 79.7% | 77.9% |
| **Avg +/- SD** | 2.57 +/- 2.83 | 2.65 +/- 3 | 2.47 +/- 2.6* | 2.72 +/- 3.02 | 2.62 +/- 3.34 |
| **95% CI** |  | 2.57 to 2.74 | 2.39 to 2.55 | 2.56 to 2.88 | 2.34 to 2.90 |
| **Median** |  | 2 | 2 | 2 | 2 |
| *Mann Whitney test, Public > Private (p = 0.026) |  |  |  |  |  |

The number of citations for first author papers was determined as a measure of the impact of the publications. F31 awardees at Private institutions were cited an average of 27.79 times per first author publication (Table 7). This was a higher citation rate than first author publications by F31 awardees from the other types of institutions, which ranged from 19.12 +/- 33.49 to 21.1 +/- 34.75 citations per paper. The number of first author publications included primary publications and reviews. The F31 awardees at Private institutions publish more reviews (9.9%) than F31 awardees at other types of institutions (6.5 to 7.7%). However, this does not account for the difference in number of citations, since F31 awardees from Private institutions have more citations for both their first author reviews and their first author primary publications (Table 7). The differences in citations for reviews is not significantly different, while the differences in citations of primary first author papers is significant.

**Table 7.** Comparison of Citations of First Author Papers of F31 Awardees.

|  | **Public** | **Private** | **Land Grant** | **IDeA** |
| --- | --- | --- | --- | --- |
| **# First Authors** | 3,453 | 2,763 | 995 | 400 |
| **# First Author Papers (All)** | 8,272 | 5,896 | 2,476 | 933 |
| **Number Citations** | 174,533 | 163,869 | 48,098 | 17,835 |
| **Avg. Citations +/- SD** | 21.10 +/- 34.75 | 27.79 +/- 46.71* | 19.43 +/- 31.86* | 19.12 +/- 33.49* |
| **95% CI** | 20.35 to 21.85 | 26.60 to 28.99 | 18.18 to 20.68 | 16.96 to 21.27 |
| **Median** | 11 | 14 | 11 | 10 |
| *Mann Whitney test, Private > Public (p < 0.0001), Land Grant < others (p < 0.0001), IDeA < others (p < 0.0001) |  |  |  |  |
| **# Reviews** | 636 | 584 | 156 | 61 |
| **% Reviews** | 7.7% | 9.9% | 6.3% | 6.5% |
| **Avg Cit./Rev +/- SD** | 36.36 +/- 49.65 | 41.91 +/- 60.51 | 34.19 +/- 43.18 | 37.64 +/- 58.24 |
| **95% CI** | 32.50 to 40.23 | 36.99 to 46.83 | 27.36 to 41.02 | 22.72 to 52.56 |
| **Median** | 21 | 23 | 20 | 17 |
| No difference |  |  |  |  |
| **# First Author Papers (excl revs)** | 7,636 | 5,312 | 2,320 | 872 |
| **Avg Cit./1<sup>st</sup> +/- SD** | 19.83 +/- 32.90 | 26.24 +/- 44.67 | 18.43 +/- 30.71 | 17.82 +/- 30.67 |
| **95% CI** | 19.09 to 20.57 | 25.04 to 27.44 | 17.18 to 19.68 | 15.78 to 19.86 |
| **Median** | 11 | 14 | 10 | 10 |
| *Mann Whitney test, Private > Public (p < 0.0001), Land Grant < others (p < 0.0001), IDeA < others (p < 0.0001) |  |  |  |  |

Analysis of the success and research productivity of F31 awardees at different types of institutions reveals comparable performance, with the exception of citations per first author publication. Given this observation, quality of students at different types of institutions does not appear to be a factor in explaining the distribution of F31 awards between different types of institutions. However, it is also possible that the success and productivity of the F31 awardees relates to the fact that they were supported by an NIH fellowship. A comparison of the publication records of postdoctoral fellows who applied for F32 fellowships, revealed increased productivity for F32 awardees compared with applicants who were not awarded an F32 (15). It is therefore of interest to compare the research productivity of a general cohort of doctoral students who graduate from different types of institutions.

#### Performance of doctoral students

Doctorates and their advisors were identified from the ProQuest Dissertations & Theses Global database. A list of doctorates publishing a dissertation in the biological and biomedical sciences from the 235 institutions designated as doctorate-granting institutions was compiled (25). Publications by the doctorates were found by searching PubMed. These searches provided lists of publications for 42,922 doctoral students completing their dissertations from 2012 to 2016 inclusive. This time frame allowed time for completion of publications after completion of training and the accumulation of citations. Further, this cohort contains the peers of a subset of the F31 cohort analyzed above. The searches captured 74,093 first author publications, 182,563 total publications.

Research productivity was measured as the number of first author publications, number of total publications and citations of first author publications. The PubMed search strategy will identify papers associated with specific doctoral students who trained with specific advisors at specific institutions, but false positives will also be included. These might be due to a shared name or might be authentic publications by the doctorate but from a different phase of their career, e.g. from undergraduate or postdoctoral studies. To partially correct for these issues, the publications chosen for analysis were constrained by date of publication and the top 1% of doctorates based upon numbers of publications were excluded as outliers.

The distribution of the differences between the publication year of papers and the year of dissertation is illustrated in Fig 10. The 5^th^ percentile of first author publications and total publications was −3, i.e. 3 years prior to the dissertation year. The 95^th^ percentile for first author publications was +4 and the 95^th^ percentile of total publications was +6. To exclude publications that were less likely to be associated with the doctoral project, i.e. publications more distant from the dissertation year, the papers selected for analysis were constrained by time. First author publications between the 5^th^ percentile (year -3) and year +2 and total publications between the 5^th^ percentile (year -3) and year +3 were included in the analyses. The upper limit is arbitrary and allows 2 years after the dissertation for publication of first author publications and 3 years after the dissertation for publication of co-authored papers. These constraints restricted the analysis to 84.1% of the first author publications from the PubMed search (62,323 papers) and 78.5% of the total publications from the PubMed search (143,381 papers). Exclusion of the doctorates greater than the 99th percentile in number of publications constrained the analysis to 42,481 doctorates (99%) with 58,602 first author publications (79.1%) and to 42,359 doctorates (98.7%) with 128,813 total publications (70.6%). Of the first author publications, 53,502 were primary peer reviewed publications and 5,100 were reviews, and 119,316 of the total publications were primary publications. The analysis of productivity was performed on these publications. Overall, students in the cohort published an average of 1.38 +/- 1.35 first author papers. Excluding published reviews, the cohort published an average of 1.26 +/- 1.25 first author papers.

**Figure 10.**
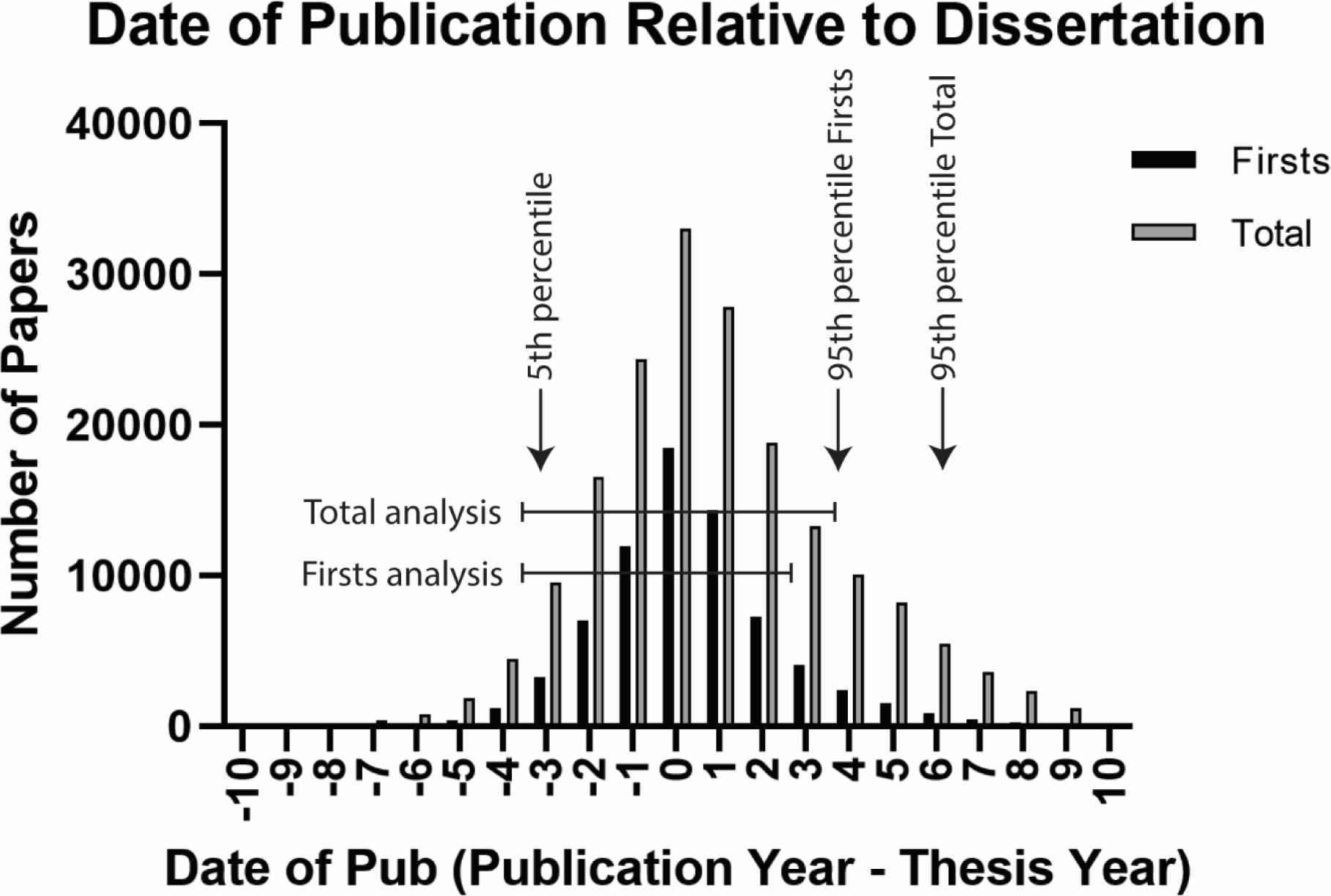
Date of publication related to dissertation year. The year of publication relative to the year of dissertation was calculated by subtracting the dissertation year from the publication year. The number of first author publications and the number of total publications in each year relative to dissertation year is plotted. The 5^th^ and 95^th^ percentiles of first author and total publications are shown. A subset of the entire dataset, restricted based on time of publication relative to the dissertation year, was used for the analysis. Lines delineate the subset of publications used for the analysis of first author papers (“Firsts analysis”) and total papers (“Total analysis”).

The productivity of students at different types of institution was measured (Table 8). A higher percentage of students at Private institutions published first author papers (70.5%) than students at Public/Land Grant institutions (66.1%/65.1%) and at IDeA institutions (61.1%). Students at Private institutions also published more reviews (12.7%) than students at Public/Land Grant institutions (9.8%/9.5%) and IDeA institutions (7.7%). The average number of first author publications, excluding reviews, was similar between students at Private, Public and Land Grant institutions (1.26 +/- 1.27 papers to 1.27 +/- 1.21 to papers). While the differences are statistically different, the differences are not meaningful and students at these institutions exhibit similar productivity by this measure. Students at IDeA institutions exhibit a lower level of productivity (1.16 +/- 1.21 papers). Given that a higher percentage of students at Private institutions publish a first author publication, it was of interest to compare the average number of publications amongst only the doctorates who have published (and excluding reviews). This analysis changes the outcome. Students at Public universities publish more first author papers on average (1.93 +/- 1.08 papers) than students at Private institutions (1.84 +/- 1.04 papers). The average first author publications of students at Land Grant universities (1.99 +/- 1.11 papers) is significantly higher than students at all other universities and the average first author publications of students at IDeA institutions (1.94 +/- 1.04 papers) is not significantly different than students at non-IDeA institutions.

**Table 8.** First Author Publications in Years -3 to +2 Inclusive (Relative to Dissertation Year)

| **All First Author Publications** |  |  |  |  |  |
| --- | --- | --- | --- | --- | --- |
|  | **ALL** | **PUBLIC** | **PRIVATE** | **LAND GRANT** | **IDeA** |
| **# Doctorates** | 42,481 | 27,907 | 14,574 | 12,760 | 4,413 |
| **# Papers** | 58,602 | 38,043 | 20,559 | 17,512 | 5,499 |
| **% Published 1st** |  | 66.1% | 70.5% | 65.1% | 61.1% |
| **Avg 1sts +/- SD** | 1.38 +/- 1.35 | 1.36 +/- 1.37 | 1.41 +/- 1.33* | 1.37 +/- 1.39* | 1.25 +/- 1.35* |
| **95% CI** |  | 1.35 to 1.38 | 1.39 to 1.43 | 1.35 to 1.40 | 1.21 to 1.29 |
| **Median** |  | 1 | 1 | 1 | 1 |
| *Mann Whitney test, Private > Public (p < 0.0001), Land Grant < others (p = 0.008), IDeA < others (p < 0.0001) |  |  |  |  |  |
| **First Author Publications Excluding Reviews** |  |  |  |  |  |
| **% Publish Rev.** |  | 9.8% | 12.7% | 9.5% | 7.7% |
| **# Papers** | 53,502 | 34,994 | 18,508 | 16,188 | 5,114 |
| **Avg 1<sup>st</sup> +/- SD** | 1.26 +/- 1.25 | 1.26 +/- 1.27 | 1.27 +/- 1.21* | 1.27 +/- 1.3 | 1.16 +/- 1.27* |
| **95% CI** |  | 1.24 to 1.27 | 1.25 to 1.29 | 1.25 to 1.29 | 1.12 to 1.20 |
| **Median** |  | 1 | 1 | 1 | 1 |
| *Mann Whitney test, Private > Public (p = 0.0005), IDeA < others (p < 0.0001) |  |  |  |  |  |
| **First Author Publications Excluding Reviews and Doctorates Who Have Not Published** |  |  |  |  |  |
| **# Doctorates** | 28,183 | 18,108 | 10,075 | 8,159 | 2,641 |
| **# Papers** | 53,502 | 34,994 | 18,508 | 16,188 | 5,114 |
| **Avg 1sts +/- SD** | 1.90 +/- 1.07 | 1.93 +/- 1.08* | 1.84 +/- 1.04 | 1.98 +/- 1.10* | 1.94 +/- 1.09 |
| **95% CI** |  | 1.92 to 1.95 | 1.82 to 1.86 | 1.96 to 2.01 | 1.90 to 1.98 |
| **Median** |  | 2 | 2 | 2 | 2 |
| *Mann Whitney test, Public > Private (p < 0.0001), Land Grant > others (p < 0.0001) |  |  |  |  |  |

A larger percentage of doctoral students at Private institutions also publish more total papers (81.3%) than students at Public/Land Grant institutions (77.2%/76%) and students at IDeA institutions (73.1%) (Table 9). Excluding reviews, students at Private institutions publish more total papers than students at other types of institutions, averaging 2.90 +/- 2.68 papers. Students at Public and Land Grant institutions publish comparable numbers of papers (2.77 +/- 2.73 papers vs 2.74 +/- 2.73 papers), while students at IDeA institutions publish fewer total papers (2.54 +/- 2.68 papers). These differences are virtually eliminated if the analysis includes only the doctorates who have published. Only students at IDeA institutions publish fewer papers on average (3.51 +/- 2.58 papers) than students at non-IDeA institutions. Based upon total publications, students at different types of institutions exhibit comparable productivity with deviations within 5% of average total publications.

**Table 9.** Total Publications in Years -3 to +3 Inclusive (Relative to Dissertation Year)

| **All Total Author Publications** |  |  |  |  |  |
| --- | --- | --- | --- | --- | --- |
|  | **ALL** | **PUBLIC** | **PRIVATE** | **LAND GRANT** | **IDeA** |
| **# Doctorates** | 42,359 | 27,823 | 14,536 | 12,733 | 4,414 |
| **# Papers** | 128,813 | 82,980 | 45,833 | 37,437 | 12,054 |
| **% Published** |  | 77.2% | 81.3% | 76.0% | 73.1% |
| **Avg Pubs +/- SD** | 3.04 +/- 2.91 | 2.98 +/- 2.92 | 3.15 +/- 2.89* | 2.94 +/- 2.92* | 2.73 +/- 2.89* |
| **95% CI** |  | 2.95 to 3.02 | 3.11 to 3.20 | 2.89 to 2.99 | 2.65 to 2.82 |
| **Median** |  | 2 | 3 | 2 | 2 |
| *Mann Whitney test, Private > Public (p < 0.0001), Land Grant < others (p < 0.0001), IDeA < others (p < 0.0001) |  |  |  |  |  |
| **All Total Author Publications Excluding Reviews** |  |  |  |  |  |
| **% Publish Rev.** |  | 16.5% | 20.0% | 16.1% | 14.5% |
| **# Papers** | 119,316 | 77,151 | 42,165 | 34,860 | 11,226 |
| **Avg Pubs +/- SD** | 2.82 +/- 2.71 | 2.77 +/- 2.73 | 2.90 +/- 2.68* | 2.74 +/- 2.73* | 2.54 +/- 2.70* |
| **95% CI** |  | 2.74 to 2.81 | 2.86 to 2.95 | 2.69 to 2.79 | 2.46 to 2.62 |
| **Median** |  | 2 | 2 | 2 | 2 |
| *Mann Whitney test, Private > Public (p < 0.0001), Land Grant < others (p < 0.0001), IDeA < others (p < 0.0001) |  |  |  |  |  |
| **First Author Publications Excluding Reviews and Doctorates Who Have Not Published** |  |  |  |  |  |
| **# Doctorates** | 33,020 | 21,304 | 11,716 | 9,577 | 3,197 |
| **# Papers** | 119,316 | 77,151 | 42,165 | 34,860 | 11,226 |
| **Avg Pubs +/- SD** | 3.61 +/- 2.56 | 3.62 +/- 2.59 | 3.60 +/- 2.52 | 3.64 +/- 2.58 | 3.51 +/- 2.58* |
| **95% CI** |  | 3.59 to 3.66 | 3.55 to 3.65 | 3.59 to 3.69 | 3.42 to 3.60 |
| **Median** |  | 3 | 3 | 3 | 3 |
| *Mann Whitney test, IDeA < others (p = 0.0006) |  |  |  |  |  |

Productivity was also measured by determining the number of citations for each first author publication, which is intended as a measure of impact in the field (Table 10). Given the difference between reviews published by students at different types of institutions, the citations of reviews and citations of publications excluding reviews were compared separately. The average number of citations of publications by students at Private institutions exceeded the citations of publications of students at other institutions. Excluding reviews, the average number of citations for doctoral students at Private institutions was 26.79 +/- 70.29 citations per paper. Students at Public and Land Grant institutions were cited 18.48 +/- 104.99 and 18.31 +/- 148.38 times per paper. IDeA institution doctoral students had an average of 13.17 +/- 18.56 citations per paper. Similarly, the average number of citations per review published by Private institution doctoral students (42.52 +/-83.15 citations per review) exceed those by students at Public, Land Grant and IDeA institutions (31.70 +/- 43.89, 29.73 +/- 36.81 and 27.25 +/- 31.98 citations per review respectively). This analysis suggests that publications by students at Private institutions has a larger impact than publications by students at other types of institutions.

**Table 10.** Citations of First Author Publications from Years -3 to +2 Inclusive (Relative to Dissertation Year)

| **Citations of All First Author Publications** |  |  |  |  |  |
| --- | --- | --- | --- | --- | --- |
|  | **ALL** | **PUBLIC** | **PRIVATE** | **LAND GRANT** | **IDeA** |
| **# Doctorates** | 28,703 | 18,437 | 10,266 | 8,307 | 2,695 |
| **# First Author Papers (All)** | 58,602 | 38,043 | 20,559 | 17,512 | 5,499 |
| **Avg Citations** | 22.64 +/- 92.36 | 19.54 +/- 101.6 | 28.37 +/- 71.87* | 19.19 +/- 143.1* | 14.17 +/- 20.14* |
| **95% CI** |  | 18.52 to 20.56 | 27.39 to 29.35 | 17.07 to 21.31 | 13.64 to 14.71 |
| **Median** |  | 10 | 14 | 10 | 9 |
| *Mann Whitney test, Private > Public (p < 0.0001), Land Grant < others (p < 0.0001), IDeA < others (p < 0.0001) |  |  |  |  |  |
| **Citations of First Author Reviews** |  |  |  |  |  |
| **# Reviews** | 5,100 | 3,049 | 2,051 | 1,324 | 385 |
| **Avg Citations** | 36.06 +/- 62.94 | 31.71 +/- 43.91 | 42.54 +/- 83.19* | 29.76 +/- 36.83* | 27.32 +/- 32.03* |
| **95% CI** |  | 30.15 to 33.27 | 38.93 to 46.14 | 27.77 to 31.74 | 24.11 to 30.53 |
| **Median** |  | 20 | 22 | 20 | 17 |
| *Mann Whitney test, Private > Public (p < 0.0001), Land Grant < others (p = 0.0016), IDeA < others (p = 0.0014) |  |  |  |  |  |
| **Citations of First Author Papers (Excluding Reviews)** |  |  |  |  |  |
| **# First Author Papers** | 53,502 | 34,994 | 18,508 | 16,188 | 5,114 |
| **Avg Citations** | 21.36 +/- 94.59 | 18.48 +/- 105.1 | 26.80 +/- 70.33* | 18.32 +/- 148.5* | 13.19 +/- 18.57* |
| **95% CI** |  | 17.38 to 19.58 | 25.79 to 27.82 | 16.03 to 20.61 | 12.68 to 13.69 |
| **Median** |  | 10 | 14 | 9 | 8 |
| *Mann Whitney test, Private > Public (p < 0.0001), Land Grant < others (p < 0.0001), IDeA < others (p < 0.0001) |  |  |  |  |  |

Another factor to consider in the analysis of scholarly activity of doctoral students is the time until the first publication. The year of matriculation into a graduate program for each student is not known, but the year of publication of the dissertation is known. Therefore, time until first publication is measured relative to the year of the dissertation using the total publication data constrained to years -3 to +3 inclusive (including papers and reviews). The first publication of students at Private institutions was -1.22 +/- 1.44 years relative to the dissertation year (Table 11). Doctoral students at Public and Land Grant institutions published their first paper -1.09 +/- 1.51 and -1.06 +/- 1.52 years relative to the year of the dissertation. Doctoral students at IDeA institutions published their first paper -0.98 +/- 1.55 years prior to the dissertation year. Students at Private institutions published earlier than students at other institutions. While the difference is significant, the difference is unlikely to make a significant real difference as Private institution students publish an average of only 1.6 to 2.9 months earlier than students at other institutions.

**Table 11.** Time to First Publication – Total Papers in Years -3 to +3 Inclusive (Relative to Dissertation Year)

|  | **All** | **Public** | **Private** | **Land Grant** | **IDeA** |
| --- | --- | --- | --- | --- | --- |
| **# Doctorates** | 42,359 | 27,823 | 14,536 | 12,733 | 4,414 |
| **# Published** | 33,290 | 21,480 | 11,810 | 9,672 | 3,226 |
| **Avg Time 1<sup>st</sup> Pub (+/- SD)** | -1.14 +/- 1.49 | -1.09 +/- 1.51 | -1.22 +/- 1.44* | -1.06 +/- 1.52* | -0.98 +/- 1.56* |
| **95% CI** |  | -1.11 to -1.07 | -1.25 to -1.20 | -1.10 to -1.03 | -1.03 to -0.92 |
| *Mann Whitney test, Private < Public (p < 0.0001), Land Grant > others (p < 0.0001), IDeA > others (p < 0.0001) |  |  |  |  |  |

These results support the conclusion that there is little meaningful difference between the average number of first author publications or the average number of total publications by students at different types of institutions. However, the first author papers published by students at Private institutions apparently have a larger impact on their field, based upon the higher number of citations per paper.

#### Comparison of the F31 cohort with the doctorate cohort

The research productivity of the F31 Awardees and the cohort of doctoral students cannot be directly compared using the NIH Reporter data and the PubMed data. To make this comparison F31 awardees were identified in the doctoral cohort by 1) matching awardee name with the name of a doctorate, 2) matching the institution where the F31 was held with the institution where the dissertation was submitted and 3) matching at least one PMID from NIH Reporter with a PMID for the PubMed search for the doctorates. Matching all these criteria resulted in the identification of 997 doctorates with first author publications who were F31 awardees and 1057 doctorates with any publication who were F31 awardees. The performance of these F31 awardees was compared with the rest of the doctorate cohort (Table 12). Since F31 awardees without publications in NIH Reporter could not be identified amongst the doctorate cohort using the criteria employed, this comparison excluded all doctorates who had not published. The F31 awardees published more first author papers (2.37 +/- 1.27 to 2.03 +/- 1.16) and more total papers (4.82 +/-3.04 to 3.84 +/- 2.74) than non F31 awardees. Citations of first author reviews by F31 awardees was comparable to the number by their peers. However, first author papers by F31 awardees, excluding reviews, receive more citations than those of their peers (27.20 +/- 49.60 to 21.12 +/- 95.96). F31 awardees also publish significantly earlier than non F31 awardees. On average the F31 Awardees publish their first paper five months earlier that the non F31 awardees. These findings suggest that F31 awardees are more productive than a large cohort of their peers.

**Table 12.** Comparison of Productivity of F31 Awardees with Their Peers.

| **First Author Publications** |  |  |
| --- | --- | --- |
|  | **Peers** | **F31 Awardees** |
| **# Doctorates** | 27,706 | 997 |
| **# First Author Papers** | 56,244 | 2,358 |
| **Avg Firsts +/- SD** | 2.03 +/- 1.16 | 2.37 +/- 1.28 |
| **95% CI** | 2.02 to 2.04 | 2.29 to 2.44 |
| **Median** | 2 | 2 |
| *Mann Whitney test, Peers < Awardees (p <0.0001) |  |  |
| **Total Publications** |  |  |
| **# Doctorates** | 32,233 | 1,057 |
| **# Total Papers** | 123,709 | 5,104 |
| **Avg Pubs +/- SD** | 3.84 +/- 2.74 | 4.83 +/- 3.04 |
| **95% CI** | 3.81 to 3.87 | 4.65 to 5.01 |
|  | 3 | 4 |
| *Mann Whitney test, Peers < Awardees (p <0.0001) |  |  |
| **Citations of First Author Reviews** |  |  |
| **# Reviews** | 4,831 | 269 |
| **Avg Citations +/- SD** | 36.01 +/- 63.85 | 37.06 +/- 43.80 |
| **95% CI** | 34.21 to 37.81 | 31.80 to 42.32 |
|  | 21 | 24 |
| *Mann Whitney test, Peers < Awardees (p = 0.15) |  |  |
| **Citations of First Author Publications (Excluding Reviews)** |  |  |
| **# First Author Papers** | 51,413 | 2,089 |
| **Avg Citations +/- SD** | 21.12 +/- 95.96 | 27.20 +/- 49.60 |
| **95% CI** | 20.29 to 21.95 | 25.07 to 29.33 |
|  | 22 | 39 |
| *Mann Whitney test, Peers < Awardees (p <0.0001) |  |  |
| **Time to First Publication** |  |  |
| **Avg +/- SD** | -1.13 +/- 1.49 | -1.56 +/- 1.29 |
| **95% CI** | -1.14 to -1.11 | -1.64 to 1.48 |
| *Mann Whitney test, Peers < Awardees (p <0.0001) |  |  |

## Discussion

The rationale for shifting support of graduate student training to fellowships and training grants is that peer review will strengthen training programs and provide some level of independence from the trainee’s research mentors (3, 4, 7). Further, there is evidence that training grants and mentored training awards are successful. Postdocs supported by F32 awards and trainees supported by K series awards exhibit higher research productivity and are more successful in the pursuit of an independent research career (11–15). The comparison of research productivity of F31 awardees versus their peers presented above is more limited in scope, but is consistent with an F31 fellowship having a positive impact on graduate student training. Given the success of these programs, expanding support would benefit a larger cohort of trainees and could qualitatively strengthen the biomedical workforce.

A striking observation about F31 funding is the divergence of the number of awardees from the number of doctorates at different types of institutions. Doctorate students from Public, Land Grant and IDeA institutions are underrepresented among F31 awardees, based upon the number of trainees at those institutions. A recent analysis of the NSF Graduate Research Fellowships Program (GRFP) and institutional factors that correlate with funding success identified affiliation with a Public institution as a factor decreasing the likelihood of success (27). The number of F31 awards at an institution correlates with the number of R01 awards at the institution, suggesting levels of NIH funding is a factor influencing F31 awards amongst institutions. Consistent with this observation, a recent analysis found that the top 14 institutions in terms of total NIH funding were also the top 14 institutions in amount of support by F series, T series and K series awards, with a very strong correlation between the top 7 in each category (28). NIH funding at an institution can affect the success of fellowship applications in several ways. It is important since it provides the resources necessary for trainees to complete their training. Further, sustained R series funding provides resources for recruitment of trainees into the lab workforce, building the training record of the PIs. Increases in the NIH budget also correlate at the macro level with increases in graduate enrollment (29, 30). The links of funding to training and training records may skew the distribution of F31 awardees to institutions with historically strong records of funding. Thus, very strong candidates in outstanding training environments may be less competitive than candidates in average training environments with very strong funding records.

The number of F31 awards received by an institution correlates with the number of doctorates awarded at the institution. Program size was also associated with funding success in the NSF GFRP (27). This may reflect a real or perceived better training program at these institutions. It may also reflect more applications from a larger doctoral pool. Data for F31 applications is not available from the NIH precluding further analysis of the applications from different types of institutions. In addition, part of the difference in number of F31 awardees at different institutions may relate to differences in the number of training grant eligible students in attendance.

### Differences in performance at different institutions

This analysis provided insight into the research productivity of doctoral students, which is of general interest in calls for transparency of outcomes of individual training programs. The results demonstrate an average of 1.38 first author publications (1.26 excluding reviews) and 3.04 total publications for a doctoral student. First author publications are cited an average of 22.64 times. The peak time for publications is in the year the dissertation is completed and the second highest number of publications appearing in the year after completion of the dissertation.

Comparison of the research productivity of F31 awardees from different types of institutions reveals little real difference in the numbers of first author publications and total publications, but papers published by F31 awardees at Private institutions are cited more often than those of other F31 awardees. Comparison of research productivity of the general population of doctoral students demonstrates that a higher percentage of students at Private institutions publish, but little real difference in the numbers of papers published. Again, papers from students at Private institutions are cited more often. The increased number of citations suggests that these papers are more impactful on the field. Quality and significance of papers are important factors in citation rates. Increased citations could reflect the higher quality of the student. Interestingly, this may also reflect the composition of the biomedical workforce at different types of institutions. The highest proportion of postdocs are in the workforce at Private institutions and interactions with postdocs as part of a cascading mentorship model of training is reported to increase the development of research skills in doctoral students (31). Citations are also impacted by extrinsic factors unrelated to the quality and significance of the paper (32). Higher citations could reflect the quality and prestige of the mentor and the institution. The number of previous publications by the author and the number of previous citations are predictive of the number of citations of future papers (32). Since doctoral students are just beginning their publication history, extrinsic factors related to citations reflect the record of the mentor, who is usually the senior author of the publication.

### Location of training and independent position

The percentage of trainees securing additional NIH funding was comparable for F31 awardees from different types of institutions. Six to eight percent of F31 awardees in the cohort secured R01 funding. This compares with reports in the literature that about 20% of F32 awardees between 2000 and 2010 also received an R01, ∼20% to 40% of K01, K08 and K23 awardees also received an R01 and ∼30% to 55% of K99 awardees also secured an R01 (11, 33). The award site for mentored career development awards (K series) impacts the distribution of trainees in their independent career since 60-80% of K01/K08/K23 awardees who receive an R01 hold their first R01 at the same institution as their K award (33). While the percentage of F31 awardees receiving R01s was not different between awardees at different institutions, the locations where the F31 awardees held their R01 awards was different. Greater than 55% of F31 awardees at Private institutions who received a subsequent R01 held their R01 at a Private institution. Seventy-four percent of F31 awardees at Public institutions who received an R01, held their R01 at a Public institution. Approximately 39% of F31 awardees at IDeA institutions held subsequent R01s at IDeA institutions. This unexpected observation suggests that the distribution of F31 awards could have an impact on where NIH-supported trainees establish their independent research careers.

### Limitations

There are a number of limitations associated with this study. Measures of research success were limited to NIH grants. Information on funding from many other federal agencies and foundations is not as robust as information from the NIH. Further, there is no mechanism to link an F31 awardee with another grant from a different agency. The emphasis on research success focused on principal investigators of subsequent grants. This is only one metric to measure research success and individuals achieving success in other ways would not be included. The publication records of F31 awardees in NIH Reporter include publications related to the award, but may not include publications prior to the award or publications related to different projects. Thus, their publication record might be incomplete. Identification of doctorates and advisors for the analysis of performance of a large cohort of doctoral students used the ProQuest Dissertations & Theses Global database. While this is a large database it is incomplete and there are institutions that do not deposit all of their dissertations in this database. Some dissertations in the dissertations database have no information on the advisor, which was required for this analysis. These records (694) were excluded. The PubMed search strategy will yield false positives when there are multiple instances of doctorates and advisors with similar names affiliated with the same institution.

### Recommendations

Institutions could take a number of actions to increase the likelihood of success of its doctoral students in winning an F31 award. Given the importance of R01 funding in the successful competition for an F31 award, institutions could create mechanisms to provide financial support to fellowship awardees and strong bridge funding commitments to their mentors, in the event of a lapse of funding. Strong commitment in support of trainees to complete their training might allay some of the fears of reviewers about potential loss of funding prior to the completion of training. Institutions could develop training programs to better prepare graduate students to compete for F31 awards, e.g. F31 writing groups (28, 34). Institutions could invest in the development, evaluation and dissemination of innovative components to their graduate curricula. Universities participating in the BEST program sponsored by the NIH set out to accomplish these goals (35). Dissemination of best practices for these types of programs is essential to impact curricula at other institutions and establish the bona fides of the training programs at institutions that currently compete poorly for F31 awards. An early time until first publication distinguishes doctorates at Private universities from doctorates at other types of universities and F31 awardees from their peers. Mechanisms created to increase the early productivity of doctoral students, in the form of co-authored publications could increase their competitiveness for F31s. This is similar to a recommendation to improve F32 competitiveness by strongly supporting early publications by postdocs (34). The other notable difference in productivity of doctorates at Private universities and F31 awardees from their peers is the number of citations of their first author papers. It is considerably more challenging to increase the impact of publications by doctoral students.

Changes in policy could also redress the skewed distribution of awardees amongst institutions. Mechanisms other than NIH grants to support student projects, including institutional commitments to support students and their mentors in the event of a lapse of funding, should be valued during fellowship review. Evaluation of the training program should emphasize innovation in training at the institution and evidence demonstrating the effectiveness of training innovations during review. This suggestion is similar to the previous recommendation to amend the peer-review process to value quality of training of all students at an institution (20). Finally, recognizing that the type of institution where an awardee holds an F31 can influence where the successful trainees hold their R01, managing the distribution of F31 awards could be an effective strategy to develop science infrastructure in underrepresented regions of the country.

## Data Availability Statement

These data were derived from the following resources at

[https://www.nsf.gov/statistics/srvygradpostdoc/, (21)],

[https://www.nsf.gov/statistics/srvydoctorates/, (22)]

## Author Contributions

MS was responsible for conceiving, designing and performing the research, data acquisition, analysis and interpretation of the data, and drafting the manuscript.

## Acknowledgments

Thanks to Dr. Marieta Gencheva for insightful comments on the manuscript. MS is the director of the Cell & Molecular Biology and Biomedical Engineering Training Program (T32 GM133369).

---

**Supplemental Figure 1.**
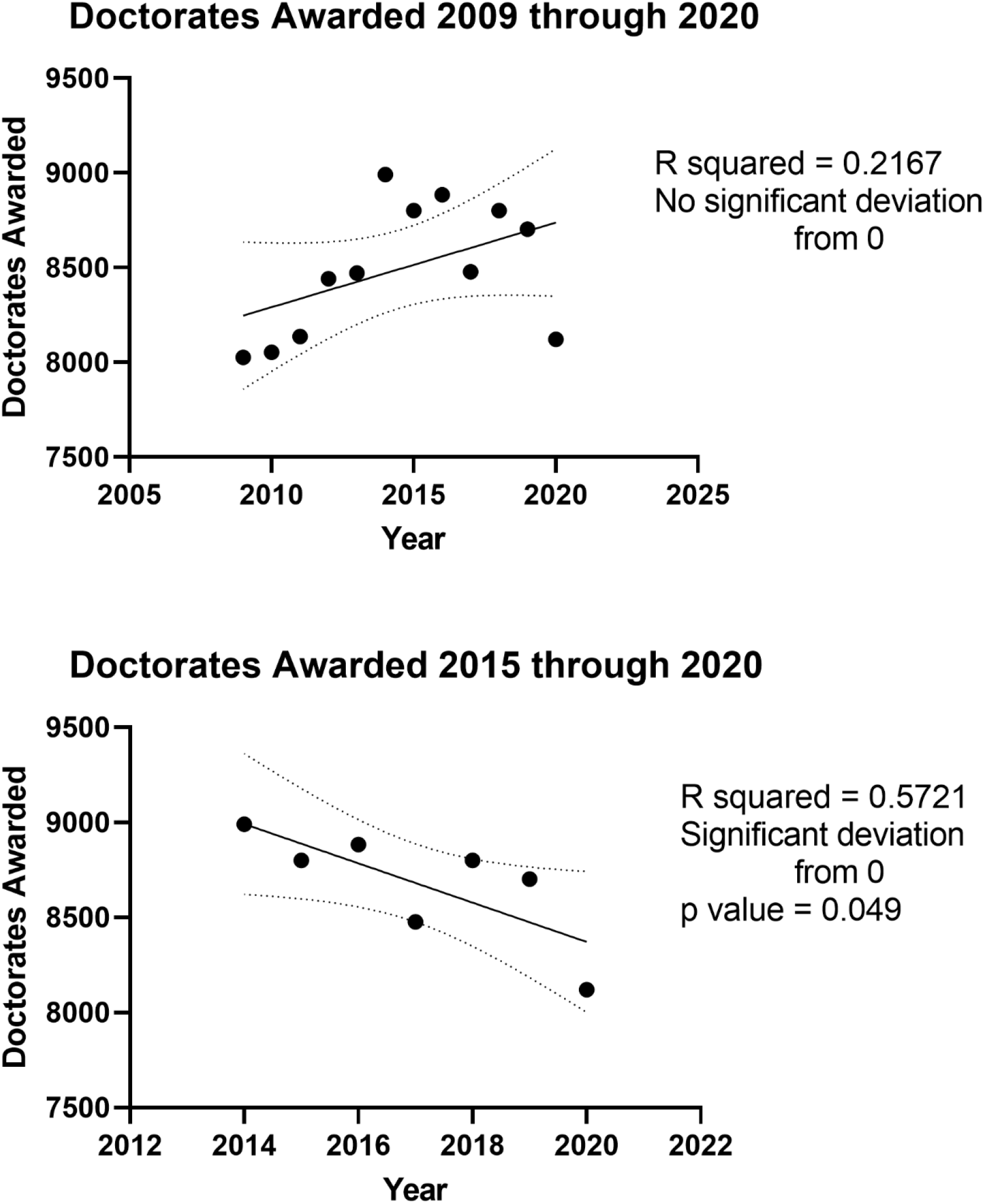
Regression analysis of number of doctorates awarded in biological and biomedical sciences. The number of doctorates awarded from 2009 through 2020 (**panel A**) and 2015 through 2020 (**panel B**) was analyzed by simple linear regression using GraphPad Prism.

